# Balanced nitrogen-iron nutrition boosts grain yield and nitrogen use efficiency in rice and wheat

**DOI:** 10.1101/2023.02.28.530550

**Authors:** Jie Wu, Ying Song, Jing-Xian Wang, Guang-Yu Wan, Zi-Sheng Zhang, Jin-Qiu Xia, Liang-Qi Sun, Jie Lu, Chuan-Xi Ma, Lin-Hui Yu, Cheng-Bin Xiang

## Abstract

Nutrients must be balanced for optimal plant growth. However, the potential significance of balanced nitrogen-iron (N-Fe) for improving crop yield and nitrogen use efficiency (NUE) has never been addressed. Here, we show that balanced N-Fe substantially increases tiller number and boosts yield and NUE in rice and wheat. NIN-like protein 4 plays a pivotal role in maintaining the N-Fe balance by coordinately regulating the expression of multiple genes in N and Fe metabolism and signaling. Moreover, we show that foliar spraying of balanced N-Fe at the tillering stage can effectively increase rice tillers, yield and NUE in the field, which can be reproduced with wheat. Our findings provide guidelines for innovative fertilization to reduce N fertilizer input and promote yield, thus benefitting environment-friendly and sustainable agriculture worldwide.

## Introduction

It is a great challenge to feed the increasing world population. To increase food production, world agriculture heavily relies on nitrogen (N) fertilizer input, whereas the fertilizer industry is energy-consuming and CO_2_-emiting (Liu et al., 2013). Reducing fertilizer input would help reduce carbon emissions. Breeding crop varieties with improved nitrogen use efficiency (NUE) is an economic and effective approach to reduce fertilizer input and increase crop yield (Liu et al., 2021; Wu et al., 2020). On the other hand, innovative fertilization technology should also be explored to enhance crop yield and minimize fertilizer input (Zhang et al., 2013).

Balanced nutrients are essential for optimal plant growth. N and iron (Fe) are widely used as fertilizers in agriculture (Briat et al., 2015; Evans and Clarke, 2019; Hu and Chu, 2020). N deficiency impairs plant growth and development and reduces crop yield (Xu and Takahashi, 2020); similarly, Fe deficiency causes chlorosis of young leaves and reduces the photosynthetic rate and respiratory intensity, ultimately leading to the disruption of energy and metabolic homeostasis (Therby-Vale et al., 2022). The balance between N and Fe represents a nutritional status that determines whether N and Fe are sufficiently abundant and optimally proportioned to maximize crop yield and NUE at a given developmental stage. Although the effects of N and Fe nutrition on plant growth have been well studied individually, the contribution of the balance between N and Fe to crop growth and yield remains largely unknown.

It is important to understand the mechanisms of nutrient balance because their interactions can produce antagonistic or synergistic outcomes that affect plant growth and yield. For example, N and phosphorus interact with each other synergistically, with nitrate promoting phosphorus utilization through the NITRATE TRANSPORTER1.1 (NRT1.1)-SPX DOMAIN-CONTAINING PROTEIN4 (SPX4) cascades (Hu and Chu, 2020). Balanced N and phosphorus supply ratios are essential for maximizing crop yields and have a greater impact on plants than sufficient N or phosphorus supplies alone (Luo et al., 2016). In contrast, it is known that phosphorus and Fe antagonize each other. A deficiency in phosphorus promotes the accumulation and utilization of Fe in plants, while excessive phosphorus causes a deficiency in Fe (Guo et al., 2022).

Interactions between N and Fe also occur in crops. For instance, the transition from N starvation to sufficient N levels results in a 3-fold increase in the Fe content in wheat shoots under stable Fe concentration conditions (Aciksoz et al., 2011). In addition, deficiencies in N and Fe cause senescence in wheat, thereby affecting the migration of Fe from its source to sinks (Parveen et al., 2018). The interaction between N and Fe has been studied to improve Fe biofortification. For example, foliar application of Fe and urea stimulates Fe absorption and translocation through wheat leaf penetration (Aciksoz et al., 2014) and significantly increases the Fe content in chickpea grains (Pal et al., 2019). However, few studies have explored the effects of the N-Fe balance on the grain yield and NUE of rice and wheat.

NIN-like protein (NLP) transcription factors are major regulators in early nitrate responses in plants (Alvarez et al., 2020). AtNLP7 responds to nitrate by accumulating in the nucleus, where the nitrate-activated calcium-dependent protein kinases (AtCPK) 10/30/32 phosphorylate AtNLP7 at Ser205, retaining AtNLP7 in the nucleus (Liu et al., 2017; Marchive et al., 2013). Recently, AtNLP7 has been shown to act as a nitrate sensor by directly binding nitrate and thus being activated (Liu et al., 2022). Our previous studies have shown that rice *OsNLPs*, as pivotal regulators of the N response, broadly regulate N assimilation-related genes and significantly increase rice yield and NUE (Alfatih et al., 2020; Wu et al., 2021; Zhang et al., 2022). Although NLPs are well known for regulating N metabolism, their roles in the regulation of the N-Fe balance have never been reported. In this study, we aimed to explore the effects of the balance between N and Fe on grain yield and NUE and revealed that balanced N-Fe significantly enhanced grain yield and NUE in an *OsNLP4*-dependent manner mainly by promoting rice tillering. Based on our novel findings, we developed a foliar fertilizer composed of N-Fe nutrients at appropriate concentrations and ratios and demonstrated that foliar application of this fertilizer at the tillering stage significantly improves yield and NUE in rice and wheat. Therefore, general guidance for improved crop fertilization can be developed based on our findings, which would have profound impacts on green and sustainable agriculture with larger yield returns and less N fertilizer input.

## Results

### Balanced N-Fe promotes rice growth and N utilization

To investigate the effect of the N-Fe balance on rice growth, we first hydroponically cultivated wild-type (WT) rice seedlings under four N-Fe conditions. Under N- and Fe-deficient (LN-LFe) conditions, the growth of the seedlings was strongly inhibited, which could be alleviated to some extent by increasing the Fe concentration. Unexpectedly, under the N-sufficient and Fe-deficient (HN-LFe) conditions, the WT seedlings exhibited the poorest growth with the most severe leaf chlorosis and produced the least amount of biomass even with sufficient N (Fig. 1a, b). Among the four N-Fe conditions, under balanced and sufficient N and Fe (HN-HFe), the WT seedlings grew the best, with the highest biomass, which was 3.2, 2.3 and 4.3 times higher than those under the LN-LFe, LN-HFe and HN-LFe conditions, respectively (Fig. 1a, b). These results were highly reproducible in multiple experimental replicates (Fig. S1) and clearly demonstrate that a balance between N and Fe tremendously impacts rice seedling growth and that HN-HFe conditions (5 mM KNO_3_:0.1 mM Fe(Ⅱ)-EDTA) are considered the appropriate N-Fe balance that results in the highest seedling biomass.

**Figure 1.**
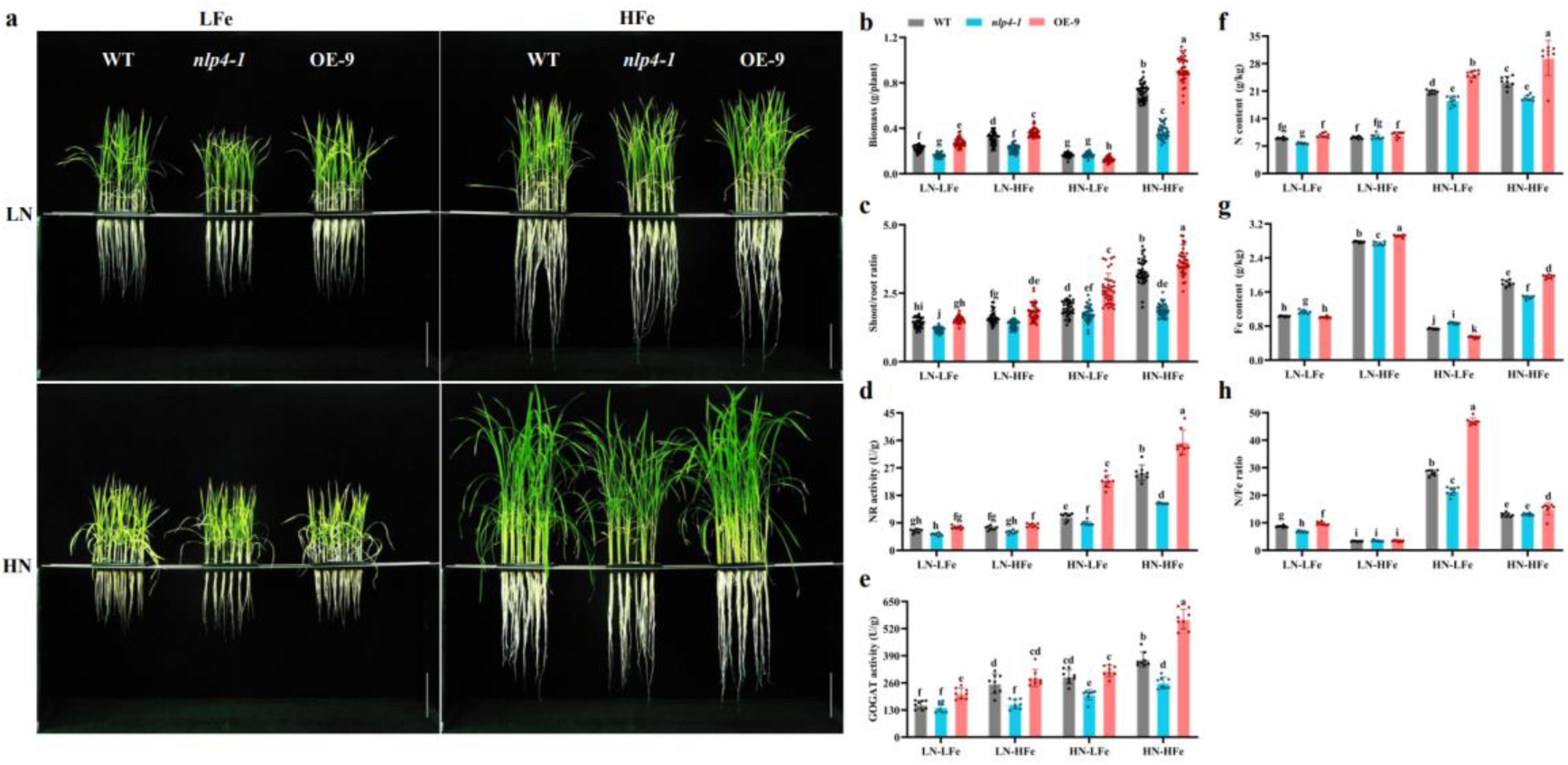
Balanced N-Fe significantly promotes the vegetative growth of rice seedlings. **a**. Phenotypes of wild-type (WT), *OsNLP4* knockout mutant (*nlp4-1*) and *OsNLP4*-OE plants (OE-9) grown in hydroponic medium with different KNO_3_ and Fe(Ⅱ)-EDTA concentrations for 28 days. LFe: 1 μM Fe(Ⅱ)-EDTA, HFe: 100 μM Fe(Ⅱ)-EDTA, LN: 0.05 mM KNO_3_, HN: 5 mM KNO_3_. Scale bar = 10.0 cm. **b-c**. Biomass (b) and shoot fresh weight/root fresh weight ratio (c) of WT, *nlp4-1* and OE-9 under different N-Fe conditions, as shown in a. **d-h**. Nitrate reductase (d) and glutamate synthase enzyme activity in leaves (e), total N content (f), Fe content (g) and total N content/total Fe content ratio (h) of WT, *nlp4-1* and OE-9 under different N-Fe conditions as shown in a. Values are the mean ± SD (n = 36 seedlings for b-c and n = 8 replicates, 6 seedlings per replicate for d-g). Different letters above bars denote significant differences (P < 0.05) from Duncan’s multiple range tests.

We then compared the *OsNLP4* knockout mutants (*nlp4-1*) and the overexpression lines (OE-9) with the WT plants under the same growth conditions. Among all the genotypes, the OE-9 plants were most responsive to the N-Fe variations, while the *nlp4* mutant plants displayed a relatively insensitive phenotype (Fig. 1a and b, S1). Under HN-HFe conditions, the OE-9 plants exhibited the best growth with a 28.3% increase in biomass compared with the WT, whereas the *nlp4* plants showed severely inhibited growth with more than 50% biomass reduction (Fig. 1b). Consistently, the shoot-to-root fresh weight ratio (S/R), nitrate reductase (NR) and glutamate synthase (GOGAT) activities displayed a similar trend as biomass in the three genotypes. The OE-9 plants exhibited significant increases in the S/R ratio, NR and GOGAT activities and total N content, which decreased in the *nlp4* mutants under HN conditions regardless of the Fe concentration (Fig. 1c-f). In addition, the OE-9 plants had the highest Fe content under HN-HFe conditions and the lowest Fe content under HN-LFe conditions (Fig. 1g), resulting in N/Fe ratios of 15.0:1 and 46.8:1 *in vivo*, which represent the most and least balanced N-Fe states, respectively (Fig. 1h). Taken together, these results demonstrate that balanced N-Fe considerably promotes rice vegetative growth and N utilization capacity and that *OsNLP4* plays a crucial role in achieving this promotion.

### Balanced N-Fe increases tillers, grain yield and NUE in rice

Encouraged by the hydroponic culture experimental results, we next assessed the effect of the balance between N and Fe on grain yield and NUE in long-term potting experiments. The grain yield of the WT plants under the four N-Fe nutrient conditions generally showed a similar trend as the seedling biomass (Fig. 2a-c). Among the N-Fe conditions, the balanced N-Fe conditions (HN-HFe) resulted in the highest grain yield, with an astonishing 10.4-fold increase in the WT compared with that under LN-LFe conditions. In contrast, under HN-LFe conditions, grain yield was the most severely compromised, with a 42-fold sharp decrease in the WT plants compared with that under HN-HFe conditions (Fig. 2b, c).

**Figure 2.**
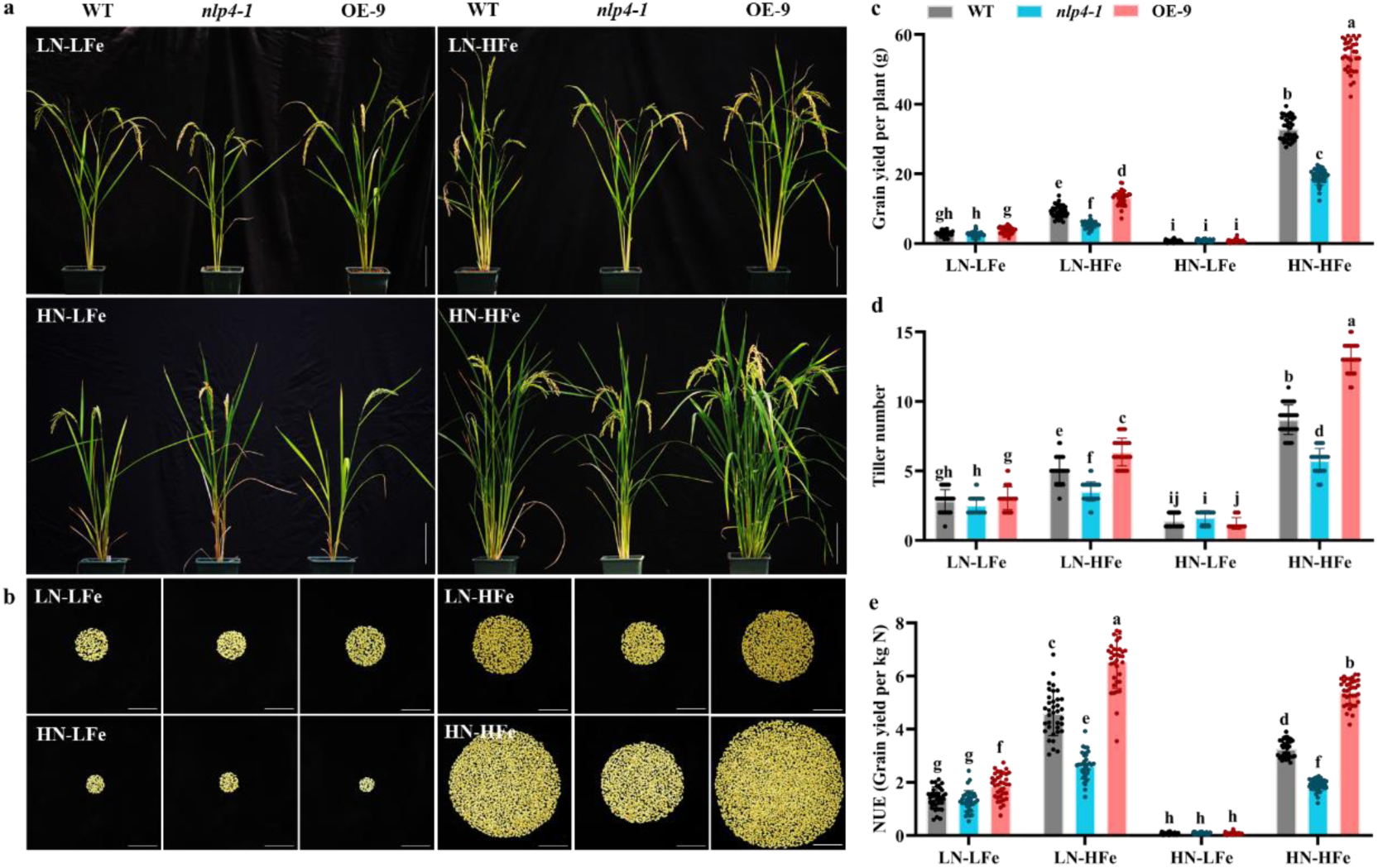
N-Fe balance dramatically affects tiller number, yield and NUE in rice. **a**. Growth phenotype at the maturing stage of representative WT, *nlp4-1*, and OE-9 grown in vermiculite pots and provided with different N-Fe nutrient solutions. Four-month-old plants were used to take images. LFe: 50 μM Fe(Ⅱ)-EDTA, HFe: 200 μM Fe(Ⅱ)-EDTA, LN: 1 mM KNO_3_, HN: 5 mM KNO_3_. Scale bar = 15.0 cm. **b**. Images of total grains per plant for WT, *nlp4-1*, and OE-9 under different N-Fe conditions as shown in a. Scale bar = 5 cm. **c-e**. Grain yield per plant (c), tiller number per plant (d) and nitrogen use efficiency (NUE) (e) of WT, *nlp4-1*, and OE-9 under different N-Fe conditions, as shown in a. Values are the mean ± SD (n = 36 plants). Different letters above bars denote significant differences (P < 0.05) from Duncan’s multiple range tests.

Among the three genotypes, the OE-9 plants were most responsive to N-Fe variations. Under HN-HFe conditions, the OE plants produced the highest yields, with a more than 65.5% grain yield increase relative to that of the WT, whereas under HN-LFe conditions, these plants produced the lowest yield, a 96.3% reduction relative to that under HN-HFe conditions. The *nlp4-1* plants produced the least amount of grain under all conditions except the HN-LFe conditions, where its yield slightly surpassed that of the OE-9 plants (Fig. 2b, c). Tiller number, a major component of yield, showed a similar change trend as grain yield in the three genotypes and was positively correlated with *OsNLP4* expression levels. Of note, under HN-HFe conditions, the tiller number of the OE-9 plants was more responsive to balanced N-Fe and substantially increased by 50.8% compared with that of the WT (Fig. 2d). Moreover, panicle length, seed setting rate, and grain weight also showed varying degrees of change under different N-Fe conditions (Fig. S2), indicating that the grain yield increase promoted by the N-Fe balance is attributed to the major contribution of tiller number supplemented by the combined effect of other yield components. These results unequivocally demonstrate that the balance between N and Fe nutrients tremendously impacts the grain yield of rice and that *OsNLP4* is a key regulator of this process.

Interestingly, NUE was positively correlated with Fe availability. Low NUE was correlated with LFe conditions regardless of the N level or genotype, with the lowest NUE occurring under the HN-LFe conditions (Fig. 2e). In contrast, high NUE was associated with high Fe availability regardless of N availability. However, NUE was slightly higher under the LN-HFe conditions than under the HN-HFe conditions, apparently due to less N input (Fig. 2e). Among the genotypes, the OE-9 plants produced the highest NUE under all N-Fe conditions except HN-LFe, where all genotypes exhibited their lowest NUE with no significant differences from each other.

Similar increases in tiller number, yield, and NUE under balanced N-Fe conditions were reproduced in multiple independent experimental replicates, as represented in Fig. S3. Taken together, our results clearly show that balanced N-Fe considerably increases tillers, grain yield, and NUE in an *OsNLP4*-dependent manner.

### Balanced N-Fe increases tillers, grain yield and NUE in wheat

Tiller number is also a major contributor to wheat yield. If balanced N-Fe promotes wheat tillering and yield as it does in rice, this outcome would be of great significance. We conducted the same potting experiments with the same N-Fe conditions for wheat as those conducted for rice. We found that an increase in either the N or Fe concentration in the nutrient solution increased the grain yield and tiller number of the wheat variety Yangmai 20. Nevertheless, compared with the HN-LFe conditions, the HN-HFe conditions resulted in a higher tiller number and grain yield, with increases of 53.7% and 90%, respectively (Fig. 3a-d). In contrast to rice, wheat experienced its worst growth under the LN-LFe conditions, where tiller number and yield decreased by 46.3% and 55.1% compared with those under the LN-HFe conditions and by 82.3% and 81.8% compared with those under the HN-HFe conditions, respectively (Fig. 3a-e). Tiller number, compared with panicle length, seed setting rate, and grain weight, showed a much stronger correlation with grain yield under different N-Fe conditions, indicating that tiller number was the major contributor to the boosted wheat yield under balanced N-Fe conditions (Fig. 3c and d, Fig. S4). A higher NUE was also positively correlated with the HFe conditions (Fig. 3e), similar to that in rice. Together, our data demonstrate that balanced N-Fe promotes tillering, yield and NUE in wheat, indicating that the underlying mechanism may be conserved in tillering gramineous plants.

**Figure 3.**
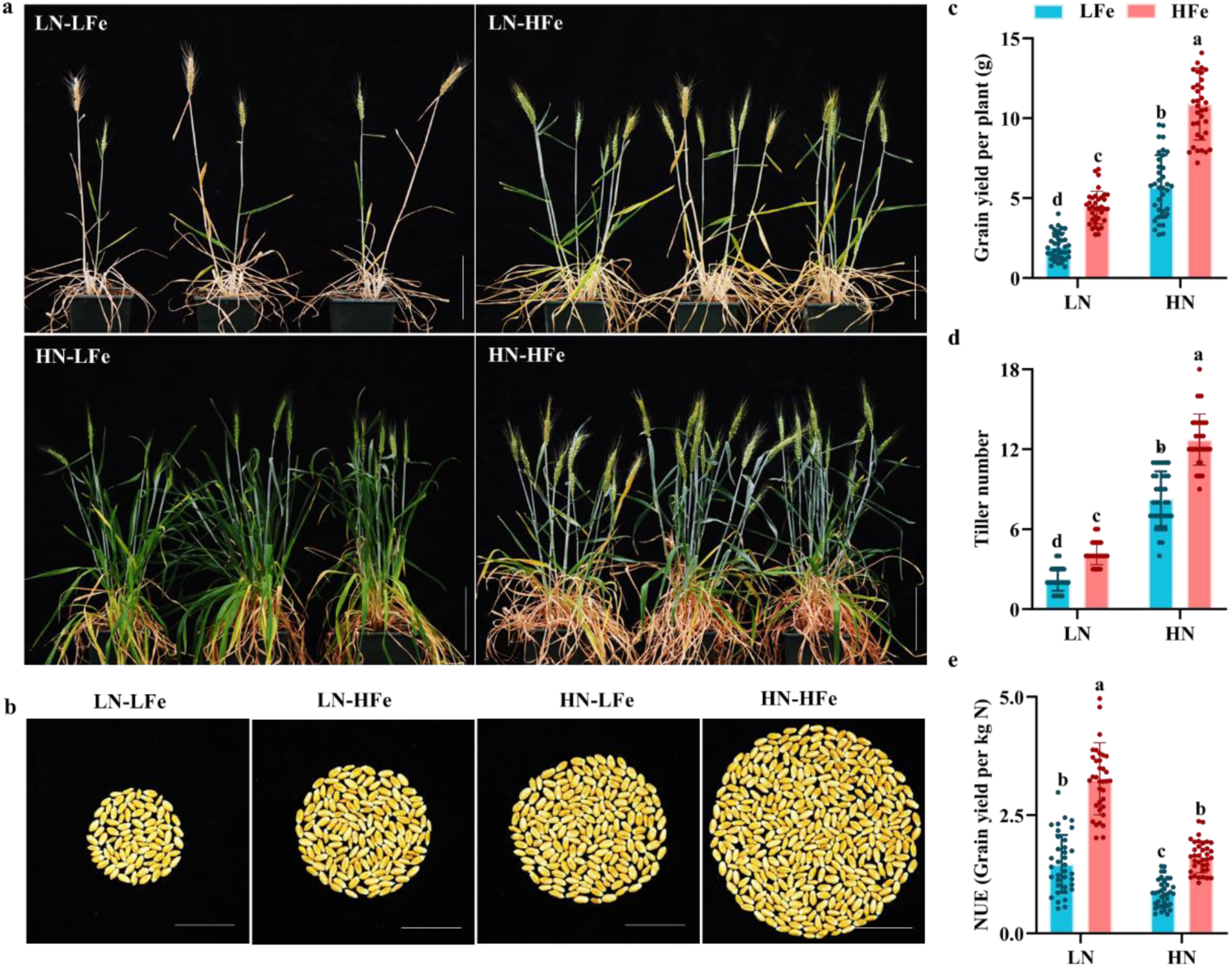
The N-Fe balance significantly affects tiller number, yield and NUE in wheat. **a**. Growth phenotype of the representative wild-type wheat cultivar Yangmai 20 (WT) at the maturation stage. Two-month-old plants were used to take images. LFe: 25 μM Fe(III)-EDTA, HFe: 150 μM Fe(III)-EDTA, LN: 1 mM KNO_3_, HN: 5 mM KNO_3_. Scale bar = 15.0 cm. **b**. Images of total grains per plant under different N-Fe conditions, as shown in (a). Scale bar = 2.5 cm. **c-d**. Grain yield per plant (c), tiller number per plant (d) and NUE (e) of the WT under different N-Fe conditions, as shown in a. Values are the mean ± SD (n = 36 plants). Different letters above bars denote significant differences (P < 0.05) from Duncan’s multiple range tests.

### Spraying of balanced N-Fe fertilizer significantly increases crop yield and NUE

To expand our findings to field crops, we first applied foliar Fe fertilizer to potted rice and wheat plants grown under HN-LFe conditions. Foliar spraying of Fe effectively restored the Fe-deficient phenotype of rice and wheat and significantly increased the number of tillers (Fig. 4a-d). Having confirmed the effectiveness of the foliar application, we next conducted field trials applying foliar Fe, N and N-Fe fertilizer to rice plants with the three *OsNLP4* genotypes during the early tillering stage under normal N conditions.

**Figure 4.**
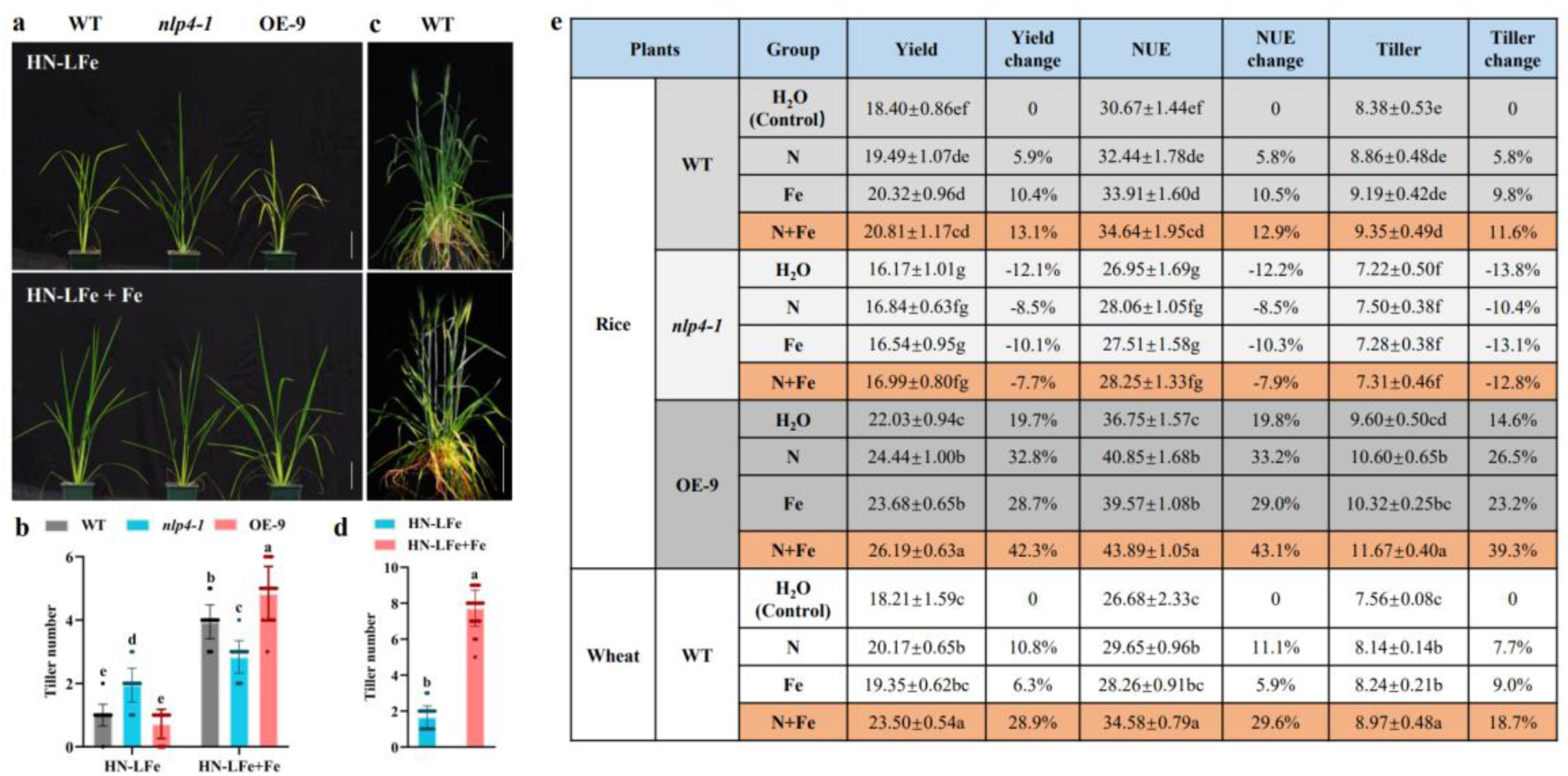
Foliar spraying of N-Fe fertilizer at the tillering stage significantly improves tillering, yield and NUE in rice and wheat in the field. **a-d**. Foliar Fe spray restores Fe-deficient growth phenotypes of rice (a) and wheat (c) plants under HN-LFe conditions. The numbers of tillers per plant of rice and wheat are shown in b and d, respectively. Plants were grown in vermiculite pots under HN-LFe conditions (top). At the early tillering stage, each plant was sprayed with 10 mL of 1 mM Fe(II)-EDTA (for rice) or Fe(III)-EDTA (for wheat) 3 times at an interval of 3 days (bottom). Two-month-old plants were used to take images. LFe: 1 μM Fe(Ⅱ)-EDTA or Fe(III)-EDTA, HN: 5 mM KNO_3_. Scale bar = 15 cm. **e**. Field trials of foliar application of N-Fe fertilizer on rice (in Lingshui, Hainan Province, China) and wheat (in Hefei, Anhui Province, China). Values are the mean ± SD (n = 18 plants for b, n = 36 plants for d, n = 4 replicates, 80 plants per replicate for rice and 40 plants per replicate for wheat for e). Different letters above bars denote significant differences (P < 0.05) from Duncan’s multiple range tests in b and e or from Student’s t test in d.

Compared with the H_2_O control treatment, the application of N and Fe fertilizer alone increased the tiller number and yield of the WT rice, whereas the N-Fe compound fertilizer markedly increased the number of tillers and grain yield by an average of 11.6% and 13.1%, respectively, resulting in an NUE enhancement of 12.9% (Fig. 4e). Compared with the WT plants treated with H_2_O, the OE-9 plants were more responsive to the N-Fe fertilizer and exhibited remarkable increases in tillers and yield of 39.3% and 42.3%, respectively, when sprayed with balanced N-Fe fertilizer (N+Fe) (Fig. 4e), whereas the *nlp4* mutant was insensitive to the foliar application of N, Fe, and N-Fe fertilizers (Fig. 4e), again confirming the important role of OsNLP4 in the response. Similar results were reproduced in another field trial at a different location (Table S1), further confirming the efficacy of our N-Fe foliar fertilizer in promoting tillering, grain yield, and NUE of rice in the field. In addition, we conducted foliar spraying tests at the heading stage and found that the foliar spray did not increase tiller number, yield or NUE even under high N conditions (Fig. S5), indicating that foliar application at the tillering stage was critical to N-Fe fertilization-promoted tillering and yield.

We also conducted a similar field trial on wheat. When N or Fe was sprayed alone, the tiller number of the wheat variety Annong 1124 showed a similar significant increase compared with the H_2_O spray control (7.7% and 9.0%, respectively), whereas the yield increased by 10.8% and 6.3%, and NUE increased by 11.1% and 5.9% (Fig. 4e). Similar to rice, when N and Fe were sprayed simultaneously, the tiller number, yield and NUE of wheat were significantly improved by 16.9%, 28.9% and 29.6%, respectively, reaffirming that the mechanism of the N-Fe balance was conserved in wheat (Fig. 4e). Taken together, these results indicate that a foliar spray application of balanced N-Fe fertilizer at the tillering stage significantly promotes rice and wheat tillering, yield and NUE in the field.

### OsNLP4 maintains the N-Fe balance in rice

To explore the role of *OsNLP4* in the regulation of the N-Fe balance in rice, we performed transcriptome analyses on seedlings cultured under different N-Fe conditions, as shown in Fig. 1a. Compared with the WT, the *nlp4* mutant and OE-9 showed very different expression profiles, with the differentially expressed genes (DEGs) varying greatly under different nutrient conditions, especially under LN-LFe and LN-HFe conditions (Fig. 5a). As shown in the Venn diagrams, very few genes are located in the overlapping parts (Fig. 5b), suggesting that OsNLP4 has a profound impact on the transcriptome of the N and Fe coresponse.

**Figure 5.**
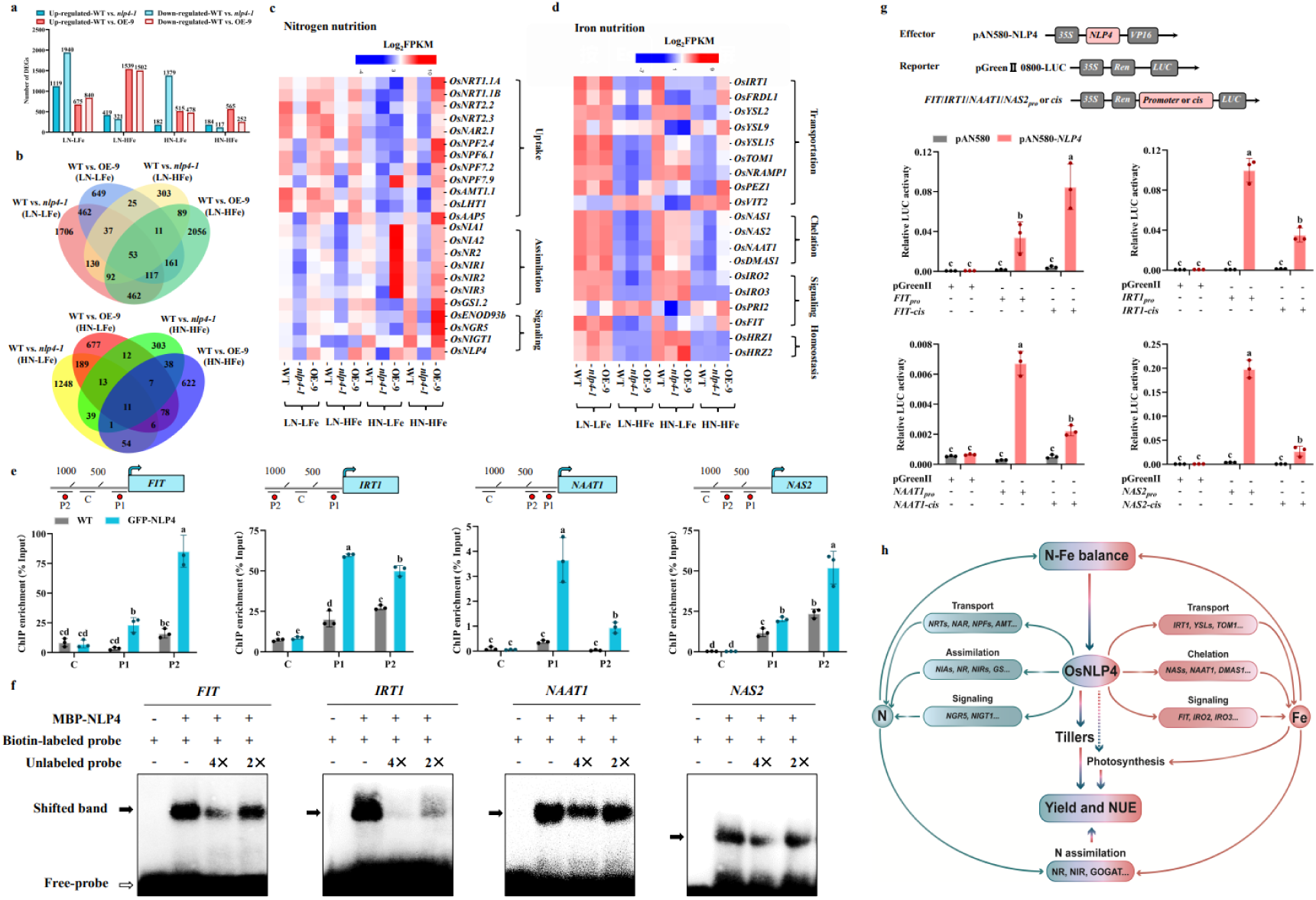
OsNLP4 regulates the coordinated expression between N and Fe nutrition-related genes. **a**. Number of upregulated and downregulated genes expressed in WT and *nlp4-1*, WT and OE-9 under different N-Fe conditions. **b**. Venn diagrams showing unique and overlapping genes significantly regulated by different N-Fe conditions. **c-d**. Hierarchical cluster analysis was performed on the expression data of 24 genes related to N nutrition (c) and 19 genes related to Fe nutrition (d) in the WT, *nlp4-1*, and OE-9 plants under different N-Fe conditions, as shown in a. The value of the blue to red gradient bar represents the log2 of the ratio. **e**. The DNA binding rate of OsNLP4 to the *OsFIT*, *OsIRT1*, *OsNAAT1* and *OsNAS2* gene promoters was analyzed by ChIP‒ qPCR. **f**. EMSA results using biotin-labeled NRE-like *cis*-fragments of *OsFIT*, *OsIRT1*, *OsNAAT1* and *OsNAS2* genes as probes and unlabeled fragments as competitors. **g**. Dual-luciferase reporting transient assays showed that OsNLP4 activates the expression of *OsFIT*, *OsIRT1*, *OsNAAT1* and *OsNAS2*. **h**. A working model of OsNLP4-modulated N-Fe balance that enhances tillering, grain yield and NUE. OsNLP4 coordinately regulates multiple genes involved in N and Fe metabolic pathways, thereby balancing N-Fe. The balanced N-Fe in turn promotes rice tillering and N assimilation in an OsNLP4-dependent manner, resulting in a significant increase in yield and NUE. In addition, Fe is required in photosynthesis and N-assimilating enzymes; thus, a sufficient Fe supply would benefit both C and N assimilation, synergistically contributing to the increased yield and NUE. Values are the mean ± SD (n = 3 replicates), and different letters above bars denote significant differences (P < 0.05) from Duncan’s multiple range tests in e and g.

We previously reported that OsNLP4 broadly regulates the expression of the genes involved in N uptake, assimilation, and signal transduction in rice (Wu *et al*., 2021). Interestingly, our transcriptomic analyses showed that the transcriptional profiles of these genes varied greatly among the different N-Fe conditions (Fig. 5c). Under HN-HFe conditions, the expression levels of these genes were significantly upregulated in OE-9 plants and sharply downregulated in *nlp4* plants compared with those in WT plants, while under HN-LFe conditions, the expression of almost all the genes involved in N absorption and signaling was unexpectedly inhibited except the N metabolism genes (*OsNIAs* and *OsNIRs*) in OE-9 plants (Fig. 5c), suggesting that Fe deficiency exerts greater impacts on N uptake and signaling than metabolism under N-rich conditions. It is worth noting that the key genes related to N uptake and assimilation were constantly downregulated in the *nlp4* plants regardless of the Fe concentration (Fig. 5c), implying a vital role of OsNLP4 in the regulation of N use by Fe.

Interestingly, we found that the transcript abundance of Fe uptake- and transport-related genes, including *OsIRT1*, *OsFRDL1*, *OsYLS2*, *OsYLS9*, *OsYLS15*, *OsYLS9*, *OsTOM1*, *OsNRAMP1*, *OsPEZ1* and *OsVIT1*, was distinctly reduced in the *nlp4* plants but significantly increased in the OE-9 plants compared with those in the WT under HN-HFe conditions (Fig. 5d). In addition to Fe transporter genes, OsNLP4 positively regulated key Fe-chelating genes, including *OsNAS1*, *OsNAS2*, *OsNAAT1* and *OsDMAS1*. Moreover, several genes involved in Fe signaling and homeostasis, such as *OsIRO2*, *OsIRO3*, *OsPRI2*, *OsFIT*, *OsHRZ1* and *OsHRZ2*, were found to be differentially expressed in the *nlp4* and OE-9 plants (Fig. 5d). Under HN-LFe conditions, the *nlp4* and OE-9 plants exhibited similar expression levels of Fe nutrition-related genes but lower levels than those of the WT (Fig. 5d). Under LN conditions, the WT and OE-9 plants showed a slight enhancement in the expression of Fe nutrition-related genes compared with the *nlp4* plants, regardless of the Fe concentration (Fig. 5d).

The expression patterns of multiple representative genes involved in N and Fe assimilation and signaling were verified by qRT‒PCR, which was generally consistent with the RNA-seq data (Fig. S6). To further study whether OsNLP4 directly regulates the expression of Fe metabolism genes, we performed *cis*-element searches within the promoters of these genes and found multiple NREs (Table S2). Subsequent chromatin immunoprecipitation (ChIP) quantitative PCR and electrophoretic mobility shift assay (EMSA) assays confirmed that OsNLP4 directly binds NRE-containing promoter fragments in Fe nutrition-related genes *in vivo*, including *OsFIT*, *OsIRT1, OsNAS2* and *OsNAAT1* (Fig. 5e-f). In addition, dual-luciferase reporter assays also demonstrated transcriptional activation of these target genes by OsNLP4 (Fig. 5g). These results indicate that OsNLP4 directly binds to the promoter of Fe nutrition-related genes and activates their expression, further supporting that OsNLP4 can regulate the expression of N- and Fe-related genes in a coordinated manner to maintain the N-Fe balance in rice (Fig. 5h).

## Discussion

N and Fe are two essential mineral elements for plant growth and development that need to remain balanced (Briat *et al*., 2015; Hu and Chu, 2020). However, the effects of the N-Fe balance on crop yield and NUE are not well understood. Here, we demonstrate that a balanced N-Fe supply promotes vegetative and reproductive growth of rice in an *OsNLP4*-dependent manner, with significantly increased NUE and yield that are mainly contributed by increased tiller number. Importantly, this balanced N-Fe-boosted yield and NUE are reproducible in wheat and can be reproduced in the field by our newly developed foliar fertilizer with balanced N-Fe.

### Balanced N-Fe significantly increases rice yield and NUE

Nutrient balance is essential for optimal plant growth and yield due to their close synergistic or antagonistic relationships in plants (Guo *et al*., 2022; Hu and Chu, 2020; Pal *et al*., 2019). Fe directly affects the activities of key enzymes in N metabolism via Fe-S clusters (Coschigano et al., 1998; Shen et al., 1976). Under N-sufficient conditions, the WT seedlings with a sufficient Fe supply significantly enhanced their N assimilation and N content (Fig. 1d-f), indicating that Fe effectively promoted rice N utilization. In addition, N participates in Fe uptake by acting as the skeleton for Fe chelates and changing the pH of the apoplast (Briat *et al*., 2015; Zou et al., 2001). The total Fe content per WT seedling under HN conditions was increased by 20% compared with that of the seedling under LN conditions when Fe was abundant (Fig. 1b, g). Therefore, balanced and sufficient N and Fe (HN-HFe) resulted in optimal rice growth and biomass and the highest yield and NUE in comparison with other conditions (Figs. 1, 2, S1, S3). Under HN-LFe conditions, rice required more Fe since N was sufficient, which exacerbated the imbalance in N and Fe (Fig. 1h) and thus resulted in worse growth than that under LN-LFe conditions (Figs. 1, 2, S1, S3). This scenario may partly explain why excessive application of N fertilizer without enough Fe did not result in a corresponding increase in the yield and NUE (Xu and Takahashi, 2020). Our study unequivocally demonstrates that balanced N and Fe significantly promote rice growth and yield in a synergistic manner and that adjusting the N/Fe supply ratio is an effective measure to increase crop productivity, which can be further optimized to reduce N fertilizer input and enhance yield.

Among the numerous grain yield determinants (Xing and Zhang, 2010), increased tillering is a major contributor to the N-Fe balance-improved rice yield and NUE. We found that tiller numbers were highly responsive to the different N-Fe conditions (Figs. 2d, 3d, 4, S3d, Table S1). It is well known that N is a key regulator of tillering in rice and wheat (Liu *et al*., 2021; Luo et al., 2020; Wu *et al*., 2020), but a direct association of Fe with tillering is rarely reported, although Fe can indirectly regulate tillering as a core component of some proteins, such as DWARF27 (Lin et al., 2009). We found that regardless of the N concentration, the number of tillers increased significantly with a sufficient supply of Fe (Figs. 2d, 3d, S3d), indicating that Fe directly promotes the occurrence of tillers. Although N and Fe alone increased tillering, the effect of the N-Fe balance was more significant. The tiller number of the WT was 5.3-fold higher under the HN-HFe conditions than under the HN-LFe conditions, resulting in an astounding 42-fold difference in yield, and wheat showed a similar difference (Figs. 2c-e, 3c-e), implying that increased tiller number is a direct cause for the N-Fe balance-boosted yield and NUE.

In this study, we uncovered a novel role of OsNLP4 in regulating tillering. The tiller number of the OE-9 plants was more responsive to different N-Fe concentrations, with a sharp 9.7-fold increase under HN-HFe conditions compared with that under HN-LFe conditions, resulting in dramatic 71- and 72-fold increases in yield and NUE, respectively (Fig. 2c-e). The *nlp4* mutant changed the least in terms of tiller number under the four N-Fe conditions (Fig. 2d), again indicating that *OsNLP4* plays a crucial role in N-Fe balance-promoted tillering in rice.

However, tillering is not the only cause for the N-Fe balance-boosted yield and NUE. Panicle length also showed a similar trend to the tillering trend among the different genotypes and N-Fe variations (Figs. S2a, S4a). In addition to increasing tillers and panicle length, balanced N-Fe also enhances N metabolism and photosynthesis because Fe is an indispensable cofactor in these processes (Briat *et al*., 2015; Coschigano *et al*., 1998; Shen *et al*., 1976), which contributes to yield and NUE.

Taken together, the balanced N-Fe enhances yield and NUE mainly through the regulation of tillering by *OsNLP4*. Transcriptome analysis indicated that OsNLP4 affects the expression of multiple genes involved in tiller development. The underlying molecular mechanisms will be reported in our companion paper.

### OsNLP4 coordinately regulates N- and Fe-related genes to balance N-Fe

NLPs are core transcription factors that regulate the early N response and N fixation (Alvarez *et al*., 2020; Jiang et al., 2021; Liu *et al*., 2022), but they have not been found to regulate Fe homeostasis and signaling pathways. Our transcriptomic data indicated that OsNLP4 broadly affected the mRNA abundance of key genes involved in Fe chelation, transport, and signaling (Fig. 5d). Biochemical assays demonstrated that OsNLP4 binds to NRE elements within the promoters of these genes to regulate their expression, suggesting that OsNLP4 is directly involved in the regulation of Fe homeostasis and signaling (Fig. 5e-g). Moreover, the Fe content increased significantly in the OE-9 plants but decreased sharply in the *nlp4* mutants under HN-HFe conditions (Fig. 1g), providing direct evidence that *OsNLP4* regulates Fe homeostasis in rice.

Interestingly, we found that OsNLP4 regulated the expression of N- and Fe-related genes in a coordinated manner, resulting in similar transcriptional profiles of both pathways in the WT, *nlp4* and OE-9 plants under the different N-Fe conditions (Figs. 5c, 5d, S6). Under HN-HFe conditions, multiple N- and Fe-related marker genes in OE-9 plants were significantly upregulated, synergistically enhancing N and Fe uptake and utilization and thus exhibiting optimal growth with the highest yield and NUE. However, under HN-LFe conditions, the expression levels of N uptake genes were more strongly inhibited in the OE-9 plants than in the WT and *nlp4* plants, which explained why the OE-9 plants lost their growth advantage (Figs. 1, 2, 5, S1, S3, Table S1). Taken together, our data suggest that OsNLP4 regulates the expression of N- and Fe-related genes in a coordinated manner, revealing the mechanism by which rice balances its own N and Fe nutrition.

### Foliar spraying of balanced N-Fe fertilizer boosts crop yield and NUE in the field

Encouragingly, the N-Fe balance-promoted agronomic traits were reproduced in the field with a foliar application of a balanced N-Fe fertilizer, which undoubtedly had great application prospects, especially for the *OsNLP4*-OE plants (Fig. 4, Table S1). A 13.1% yield increase was obtained with an Fe input at an almost negligible cost for WT rice. If OE-9 plants are planted, then grain yield will increase significantly to 42.3%. While obtaining considerable economic profits, an improvement in NUE can also reduce the environmental damage caused by excessive application of N fertilizer, which is conducive to green and sustainable agriculture (Xu and Takahashi, 2020; Zhang et al., 2015). Based on a 12.9% increase in NUE, our balanced N-Fe foliar spraying is estimated to reduce N fertilizer in WT paddy fields by 11.5% compared to conventional fertilization for the same yield and by 30% if combined with the *OsNLP4*-OE plants (Fig. 4, Table S1). It is conceivable that the input of N fertilizer can be further reduced by optimizing the N-Fe fertilizer and its application. In addition, the N-Fe balance-promoted grain yield and NUE were reproduced in another cereal crop, wheat (Figs. 3, 4), suggesting that the underlying mechanisms are conserved in gramineous plants and illuminating a potential wide application of our balanced N-Fe foliar fertilizer. It should be noted that the timing of the foliar application of the N-Fe fertilizer is critical, and this application is effective during the tillering stage but not during the heading stage (Fig. S5). The contribution of the balanced N-Fe fertilizer to crop yield and NUE in the field will provide guidance to formulate innovative fertilizers for greener agriculture.

Overall, we reveal that the N-Fe balance greatly promotes tiller number in rice and wheat, resulting in a remarkable increase in NUE and yield, while OsNLP4 balances N and Fe by regulating N- and Fe-related genes in a coordinated manner (Fig. 5h). Our balanced N-Fe foliar fertilizer can easily reproduce the N-Fe balance-promoted yield and NUE in the field, achieving economic benefits while avoiding excessive application of N fertilizer. Moreover, the conserved mechanism of N-Fe balance-promoted tillering implies that this novel finding is applicable to all crops with tillering, which would benefit diet-diverse populations globally.

## Methods

### Plant materials

All rice (*Oryza sativa)* plants used in this study were derived from the japonica cultivar Zhonghua 11 (ZH11). The *OsNLP4* knockout mutant and overexpressing plants were previously used lines (Wu *et al*., 2021). The mutant *osnlp4-1* was acquired from Bogle Hangzhou Co., Ltd. (Hangzhou, China) (http://www.biogle.cn/) using CRISPR/Cas9 technology. The homozygous overexpression line OE-9 (T4 generation) was obtained by *Agrobacterium tumefaciens* (EHA105) transformation and glufosinate screening. The wheat varieties Yangmai 13 and Annong 1124 used in this study were from Professor Chuanxi Ma of Anhui Agricultural University.

### Plant growth conditions

#### Seedlings in hydroponic culture

Seeds of WT, *nlp4-1*, and OE-9 were washed with distilled water and incubated at 37 °C for 3 days. Germinated seeds were transferred to modified Kimura B solution (pH 5.8) with different N and Fe availability treatments (LN: 0.05 mM KNO_3_, HN: 5 mM KNO_3_, LFe: 1 μM Fe(Ⅱ)-EDTA, HFe: 100 μM Fe(Ⅱ)-EDTA) to grow for 28 days. Nutrient solutions were replaced every two days. The provided growth conditions were kept at a 16-h light (30 °C)/8-h dark (28 °C) cycle in a climate chamber.

#### Seedlings in vermiculite for long-term treatment

Seeds of WT, *nlp4-1*, and OE-9 were germinated and then transferred to pots filled with vermiculite (the pot dimensions were 15×15×15 cm^3^). A single plant was grown per pot. Each treatment contained 50 pots for each genotype. Plants were fed different N-Fe solutions (LN: 1 mM KNO_3_, HN: 5 mM KNO_3_, LFe: 50 μM Fe(Ⅱ)-EDTA, HFe: 200 μM Fe(Ⅱ)-EDTA) until maturity. Each treatment was irrigated with 10 L of nutrient solution each time. During the whole growth period, plants were irrigated 12 times at intervals of 12 days. For wheat, plants were fed different N-Fe solutions (LN: 1 mM KNO_3_, HN: 5 mM KNO_3_, LFe: 25 μM Fe(III)-EDTA, HFe: 150 μM Fe(III)-EDTA) until maturity. Each treatment was irrigated with 10 L of nutrient solution each time. During the whole growth period, plants were irrigated 8 times at intervals of 15 days. The provided growth conditions were kept at a 12-h light (30 °C)/12-h dark (28 °C) cycle in a greenhouse. NUE was defined as the grain yield per unit of available N (KNO_3_) in the vermiculite (Hawkesford, 2014).

### Enzyme activity assay and metabolite analyses

Twenty-eight-day-old hydroponic seedlings were used for all assays in this section. NR and GOGAT activities were analyzed with an enzyme-coupled spectrophotometer assay kit (SKBC, China, cat # AKNM1001M and AKAM009M) according to the manufacturer’s guidelines. For the total N content assay, whole seedlings were dried at 70 °C to a constant weight, ground and then analyzed with an NC analyzer (Vario EL III model, Elementar, Hanau, Germany) according to the manufacturer’s instructions. Leaf chlorophyll was extracted using 80% acetone and measured by a spectrophotometric method as described previously (Wu *et al*., 2021). For the total Fe content assay, whole seedlings or seeds were dried at 70 °C to a constant weight, ground, and then wet-ashed with HNO_3_, perchlorate and H_2_O_2_ for 1∼2 hours at 220 °C using a nitrate boiling tube. The Fe concentration was measured by inductively coupled plasma–mass spectrometry (ICP–MS) as previously described (Guo *et al*., 2022).

### RNA-seq analyses

Each strain (approximately 96 seedlings, ZH11 background) with different N-Fe (0.05/5 mM KNO_3_ and 1/100 μM Fe(Ⅱ)-EDTA) treatments was hydroponically grown in an artificial climate chamber under the conditions described above. Two-week-old seedlings (whole plant) were selected for RNA sequencing. Thirty seedlings were collected from each treatment as samples for three independent biological replicates. RNA library construction and sequence analysis were described previously (Wu *et al*., 2021).

### qRT‒PCR analyses

Total RNA was isolated from 2-week-old rice seedlings using TRIzol reagent (TransGen, cat # ET111). Full-length cDNA was then reverse transcribed using a cDNA synthesis kit (TRANSGEN, cat # AE311-02). The qRT‒PCR step one Plus real-time PCR system was used for the TaKaRa SYBR premix Ex Taq II reagent kit (cat # Q111-02). Subsequent qRT‒PCR procedures were performed according to the manufacturer’s instructions, and each qRT‒PCR assay was repeated at least three times with three independent RNA preparations. Rice Actin1 was used as an internal reference. The primers used are shown in Table S3.

### ChIP-PCR assay

After 7 days of growth in high-Fe rice nutrient solution (100 μM), the seedlings were transferred to low-iron rice nutrient solution (1 μM) for 7 days, and then the whole seedlings were taken for the ChIP experiment. Samples (3-4 g) of *OsACTIN1pro:OsNLP4-GFP* transgenic plants (whole seedlings) were fixed and crosslinked with 1% formaldehyde vacuum for 10 min, and then 125 mM glycine was added to stop the reaction. The samples were frozen in liquid nitrogen and then ground to a powder. Ultrasound at low temperature (4 ℃) fragmented chromatin to an average size of 400-600 bp. GFP-trap magnetic beads are used for immunoprecipitation of protein DNA complexes. Chromatin without antibody precipitation was used as an internal reference, and the gene enrichment level was detected by real-time PCR.

### Protoplast transfection and dual-luciferase reporter assay

Stems from rice growing at 2-3 weeks old were used to prepare protoplasts. In protoplast transient expression experiments, plasmids were transfected into protoplasts as described previously (Liu *et al*., 2021). One thousand bp upstream of the target gene initiation codon and the NRE-like element (*cis*-fragment) were cloned into the vector pGreen II0800-LUC to generate the reporter gene for dual-luciferase detection. The full-length *OsNLP4* CDS is inserted into Pan580 to generate the effector. Firefly luciferase (LUC) activity and Renilla luciferase (REN) activity were determined using a dual luciferase reporting system (Vazyme, cat # DL101-01) 12-18 h after transfection. The LUC and REN activity ratios were determined at least three times.

### EMSAs

EMSAs were performed as described previously (Liu et al., 2018). Purified recombinant MBP-OsNLP4 protein and GST protein from Escherichia coli. *Cis*-fragments in promoters of potential target genes of OsNLP4 were synthesized, and biotin was labeled at the DNA ends. Unlabeled fragments of biotin with the same sequence or mutant sequence were used as competitors, and MBP protein was used as a negative control. The DNA gel shift was measured using the LightShift Chemiluminescence EMSA Kit (Thermo Fisher Scientific, cat # 20148).

### Field trials for foliar application of N-Fe fertilizer

#### Rice field trial

The field foliar spray experiment comprised three *OsNLP4* genotypes (wild type, knockout mutant *nlp4-1*, and overexpression line OE-9) and four treatments: i) control (foliar application of H_2_O); ii) foliar application of N (0.5% urea); iii) foliar application of Fe (1 mM Fe(Ⅱ)-EDTA); and iv) simultaneous foliar spray of Fe (1 mM Fe(Ⅱ)-EDTA) in a cocktail solution containing 0.5% urea. At the tillering or heading stage, plants received foliar spray 3 times at an interval of 5 days. All foliar solutions were applied at 1600 liters/hectare each time. The foliar treatments were applied in the late sunny afternoon as described previously (Prom et al., 2020). All treatments received the same basic management. The field experiment was laid out in a randomized complete block design with four replications. Field tests for rice were carried out in paddy fields under natural growth conditions during 2020-2022 at three experimental locations: Bengbu (Anhui Province, China), Lingshui (Hainan Province, China), and Changxing (Zhejiang Province, China).

The Bengbu field trial was performed in 2020 (May to October). Urea was used as the N source at 185 kg N/ha. The plants were transplanted in 8 rows×10 plants for each plot (3.6 m^2^), and four replicates were used for each treatment.

The Lingshui field test was performed from December 2021 to April 2022, and urea was used as the N source at 150 kg N/ha. The plants were transplanted in 10 rows×10 plants for each plot (4 m^2^), and four replicates were used for each treatment.

The Changxing field test was conducted in 2021 (May to October). Urea was used as the N source, with 180 kg N/ha for normal N and 360 kg N/ha for high N. The planting density was 8 rows×10 plants for each plot (3.4 m^2^) with four replicates.

#### Wheat field trial

All foliar spray treatments were the same as those for rice except Fe(II)-EDTA was replaced with Fe(III)-EDTA. Field trials for the wild-type wheat cultivar Annong 1124 were carried out at the experimental station of Anhui Agricultural University in Hefei from November 2021 to May 2022. Urea was used as the N source at 270 kg N/ha. The plants were transplanted in 20 rows×10 plants for each plot (5 m^2^), and four replicates were used for each treatment.

For data collection, the edge lines of each plot were excluded to avoid margin effects. NUE was defined as the grain yield per unit of available N (urea) in the soil (Hawkesford, 2014).

### Statistical analysis

The data were statistically analyzed, and multiple comparisons were performed using Duncan’s multiple range tests. A P value less than 0.05 was considered statistically significant.

### Data and materials availability

The data supporting the findings of this study are available within this paper and its Supplementary information files. The sequence data used in this study can be found in the Rice Genome Annotation Project (https://rice.plantbiology.msu.edu/) under the following accession numbers: *OsNLP4*, *LOC_Os09g37710*; *OsNIA1*, *LOC_Os08g36480*; *OsNR2*, *LOC_Os02g53130*; *OsNiR1*, *LOC_Os01g25484*; *OsFIT*, *LOC_Os04g31290*; *OsIRT1*, *LOC_Os03g46470*; *OsNAAT1*, *LOC_Os02g20360*; *OsNAS2*, *LOC_Os03g19420*. The datasets and materials used during the current study are available from the corresponding author on reasonable request.

## Author Contributions

J.W. and C.B.X. designed the experiments. J.W., Y.S., J.X.W. and G.Y.W. performed the experiments and data analyses. J.W., Z.S.Z., J.Q.X., L.Q.S., J.L. and C.X.M. performed field trials and data analyses. J.W. wrote the manuscript. C.B.X., L.H.Y., and J.W. revised the manuscript. C.B.X. supervised the project.

## Acknowledgments

We thank Drs. Chuan-Zao Mao (College of Life Sciences, Zhejiang University) and Shi-Mei Wang (Rice Research Institute, Anhui Academy of Agricultural Science) for their assistance with field trials, Dr. Chengcai Chu (Institute of Genetics and Developmental Biology, CAS, Beijing, China) for his critical comments and suggestions on the manuscript, and Dr. Chunming Wang (Nanjing Agricultural University, Nanjing, China) for providing the pAN580 plasmid.

---

**Figure S1.**
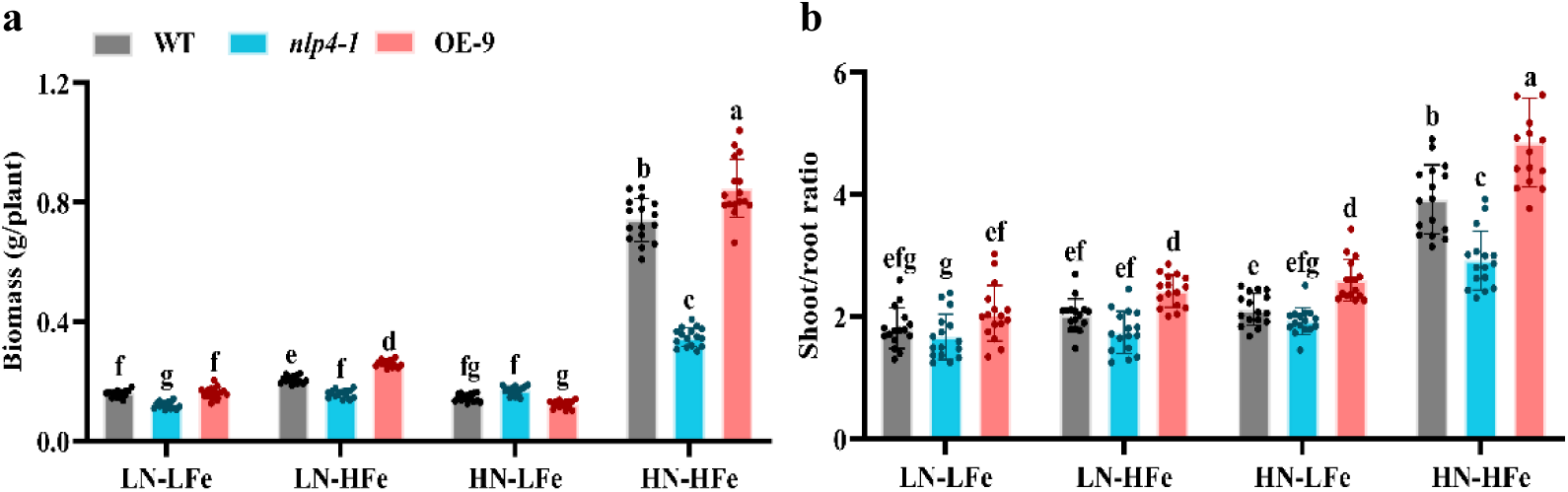
The N-Fe balance dramatically affects the vegetative growth of rice seedlings. **a-b**. Biomass (a) and shoot fresh weight/root fresh weight ratio (b) of wild type (WT), *osnlp4* mutant (*nlp4-1*) and *OsNLP4*-OE plants (OE-9) grown in hydroponic medium with different KNO_3_ and Fe(Ⅱ)-EDTA concentrations. Germinated seeds were transferred to modified Kimura B solution (pH 5.8) with different N and Fe availability treatments (LN: 0.05 mM KNO_3_, HN: 5 mM KNO_3_, LFe: 1 μM Fe(Ⅱ)-EDTA, HFe: 100 μM Fe(Ⅱ)-EDTA) to grow for 28 days. Nutrient solutions were replaced every two days. Biomass represents the fresh weight of the whole seedling. Values are the mean ± SD (n = 16 seedlings). Different letters above bars denote significant differences (P < 0.05) from Duncan’s multiple range tests.

**Figure S2.**
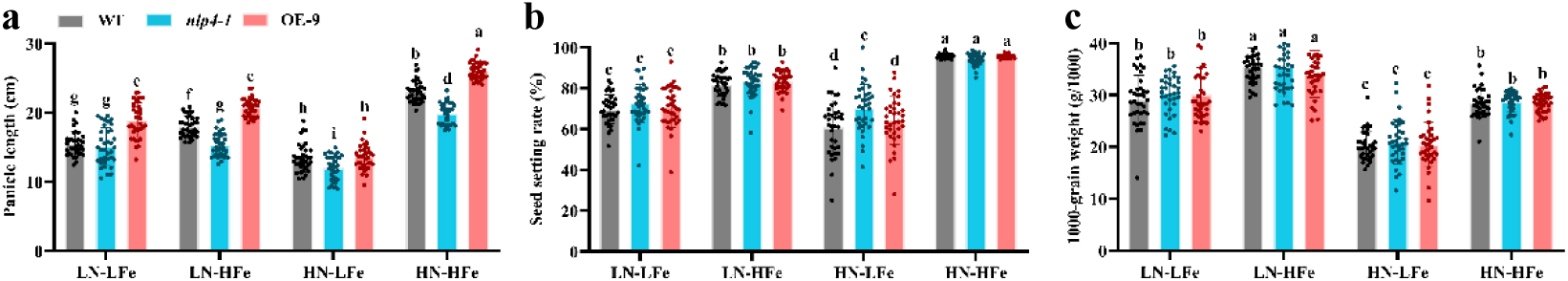
Agronomic traits of rice under different N-Fe regimes. **a-c**. Panicle length (a), seed setting rate (b), and 1000-grain weight (c) of WT, *nlp4-1* and OE-9 under different N-Fe treatments. Seeds were germinated and then transferred to pots filled with vermiculite. A single plant was grown per pot. Every treatment contained 50 pots for each genotype. Each treatment was irrigated with 10 L of nutrient solution each time. During the whole growth period, plants were irrigated 12 times at intervals of 12 days. LFe: 50 μM Fe(Ⅱ)-EDTA, HFe: 200 μM Fe(Ⅱ)-EDTA, LN: 1 mM KNO_3_, HN: 5 mM KNO_3_. Values are the mean ± SD (n = 36 plants). Different letters above bars denote significant differences (P < 0.05) from Duncan’s multiple range tests.

**Figure S3.**
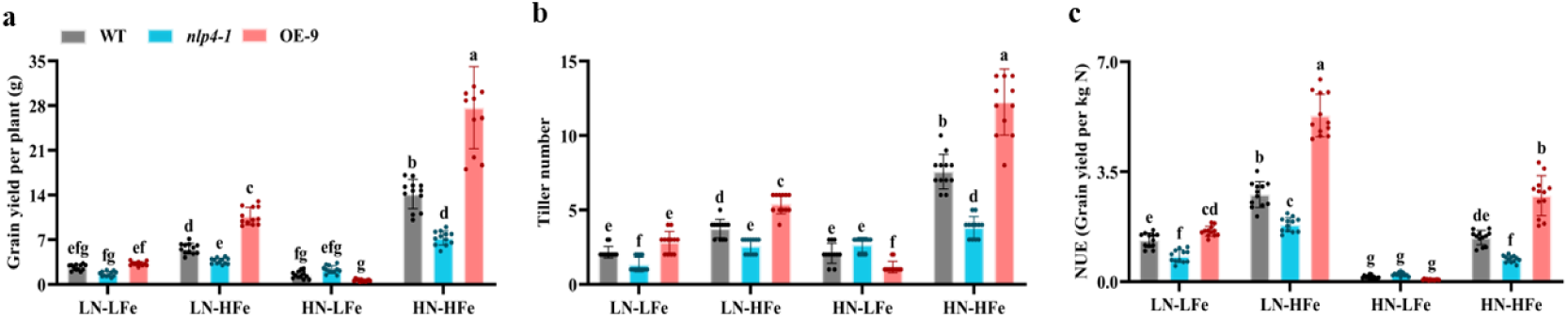
Different N-Fe regimes greatly influence tiller number, yield and NUE in rice. **a-c**. Grain yield per plant (c), tiller numbers per plant (d) and nitrogen use efficiency (NUE) (e) of WT, *nlp4-1*, and OE-9 under different N-Fe treatments. Seeds were germinated and then transferred to pots filled with vermiculite. A single plant was grown per pot. Every treatment contained 50 pots for each genotype. Each treatment was irrigated with 10 L of nutrient solution each time. During the whole growth period, plants were irrigated 12 times at intervals of 12 days. LFe: 50 μM Fe(Ⅱ)-EDTA, HFe: 200 μM Fe(Ⅱ)-EDTA, LN: 1 mM KNO_3_, HN: 5 mM KNO_3_. Values are the mean ± SD (n = 12 plants). Different letters above bars denote significant differences (P < 0.05) from Duncan’s multiple range tests.

**Figure S4.**
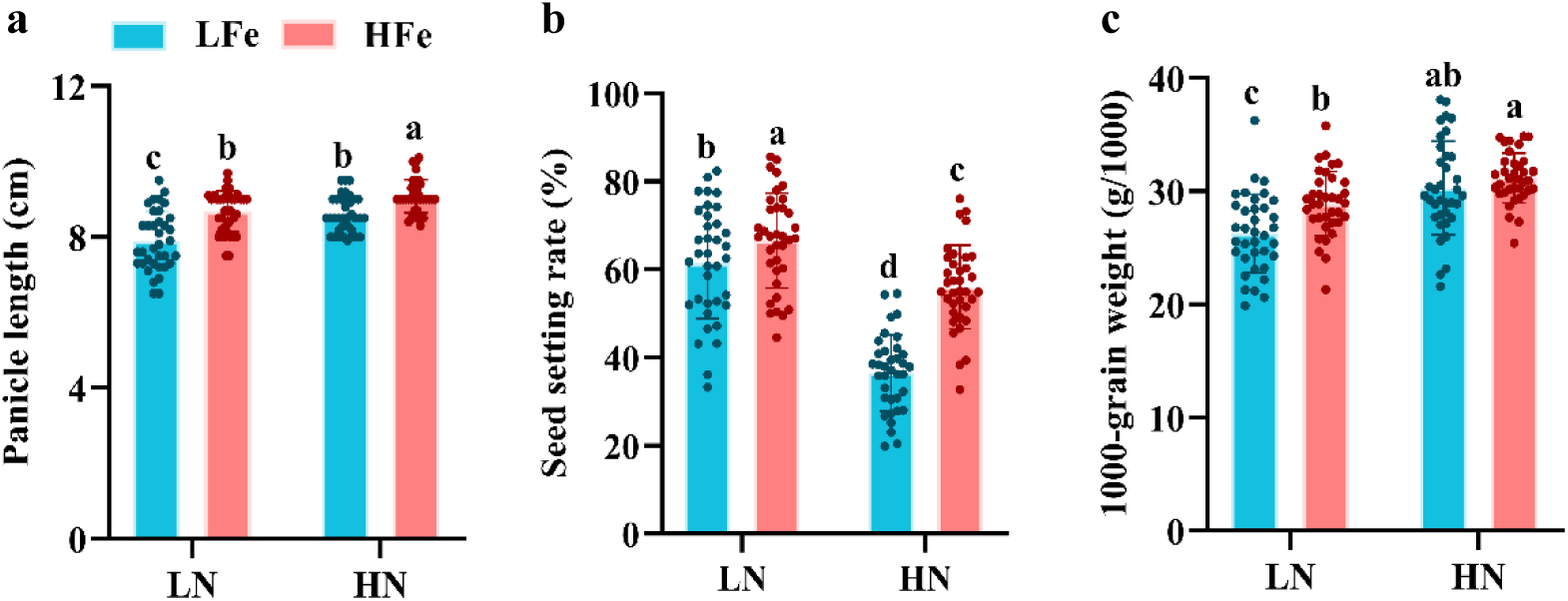
Agronomic traits of wheat under different N-Fe regimes. **a-c**. Panicle length (a), seed setting rate (b) and 1000-grain weight (c) of wheat under different N-Fe treatments. Seeds were germinated and then transferred to pots filled with vermiculite. A single plant was grown per pot. Every treatment contained 50 pots. Each treatment was irrigated with 10 L of nutrient solution each time. During the whole growth period, plants were irrigated 8 times at intervals of 15 days. LFe: 25 μM Fe(III)-EDTA, HFe: 150 μM Fe(III)-EDTA, LN: 1 mM KNO_3_, HN: 5 mM KNO_3_. Values are the mean ± SD (n = 36 plants). Different letters above bars denote significant differences (P < 0.05) from Duncan’s multiple range tests.

**Figure S5.**
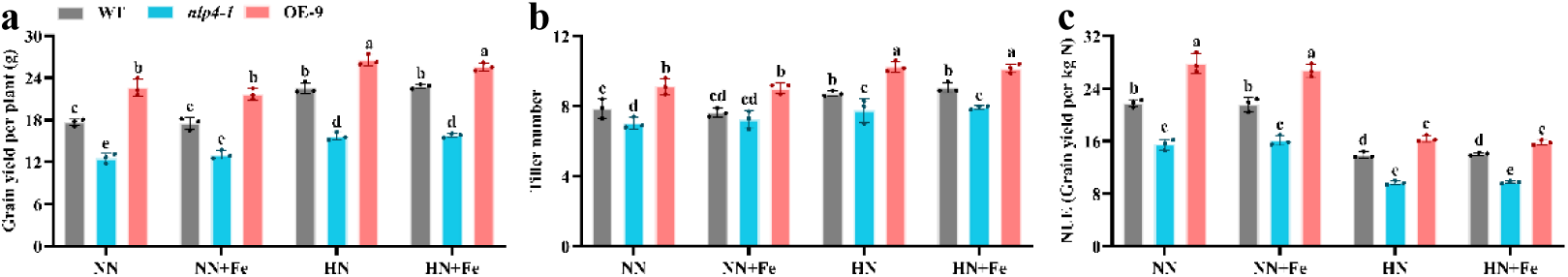
Foliar spraying of Fe microfertilizer at the heading stage did not contribute to rice yield and NUE. **a-c**. Grain yield per plant (a), nitrogen use efficiency (NUE) (b) and tiller numbers per plant (c) of WT, *osnlp4* mutants (*nlp4-1*) and *OsNLP4*-OE plants (OE-9) sprayed with Fe fertilizer under different N concentrations at the heading stage in Changxing in 2021 (May to October). Urea was used as the N source, with 180 kg N/ha for normal N and 360 kg N/ha for high N, and the planting density was 8 rows×10 plants for each plot (3.4 m^2^) with four replicates. NN, normal N conditions. HN, high N conditions. At the early heading stage, each plant was sprayed with 1 mM Fe(Ⅱ)-EDTA 10 mL 3 times, with an interval of 5 days. Values are the mean ± SD (n = 3 replicates, 60 plants per replicate). Different letters above bars denote significant differences (P < 0.05) from Duncan’s multiple range tests.

**Figure S6.**
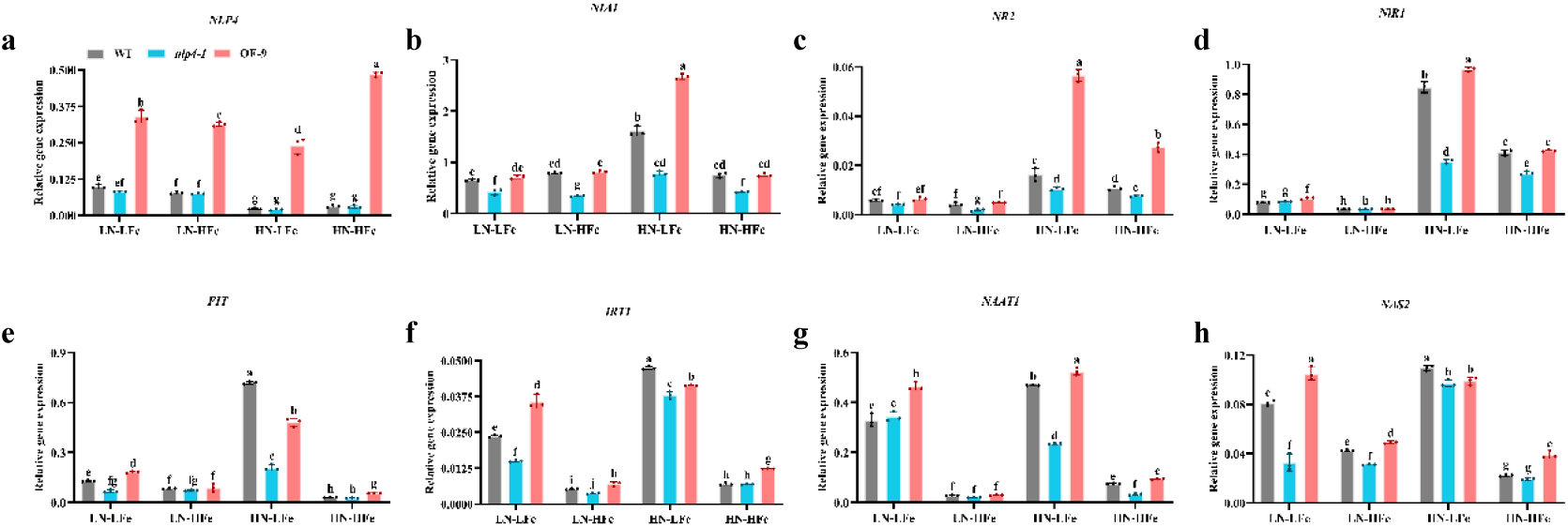
Transcription levels of selected genes involved in N and Fe metabolism and signaling pathways revealed by qRT‒PCR. Twenty-eight-day-old seedlings (whole plants) cultured with different N-Fe concentrations (LFe: 1 μM Fe(Ⅱ)-EDTA, HFe: 100 μM Fe(Ⅱ)-EDTA, LN: 0.05 mM KNO_3_, HN: 5 mM KNO_3_) were sampled for RNA. *OsActin1* was used as an internal reference. qRT‒PCR data are the mean ± SD (n = 3 replicates). Different letters above bars denote significant differences (P < 0.05) from Duncan’s multiple range tests. *NLP4*: NIN-like protein 4, *NIA* and *NR*: nitrate reductase, *NiR1*: nitrite reductase 1, *FIT*: fer-like Fe deficiency-induced transcription factor, *IRT1*: iron-regulated metal transporter, *NAAT1*: nicotianamine aminotransferase *1*, *NAS2*: nicotianamine synthase.

**Table S1.** Foliar spraying of N-Fe compound fertilizer at the tillering stage significantly improves rice tillering, yield and NUE in the field.

| Plant | Group | Yield | Yield change | NUE | NUE change | Tiller | Tiller change |
| --- | --- | --- | --- | --- | --- | --- | --- |
| WT | H<sub>2</sub>O (Control) | 20.69±1.46f | 0 | 25.04±1.02f | 0 | 9.66±0.35ef | 0 |
|  | N | 22.28±1.11de | 7.6% | 26.88±1.61e | 7.4% | 10.29±0.31de | 5.8% |
|  | Fe | 22.02±1.12e | 6.4% | 26.73±1.30e | 6.8% | 10.22±0.24de | 9.8% |
|  | N+Fe | 23.24±1.24d | 12.3% | 28.31±1.44d | 13.0% | 10.63±0.27cd | 11.6% |
| *nlp4-1* | H<sub>2</sub>O | 17.69±0.96g | -14.5% | 21.22±1.10g | -15.2% | 8.62±0.17g | -10.1% |
|  | N | 18.23±0.80g | -11.9% | 21.87±1.24fg | -12.7% | 8.83±0.23fg | -8.5% |
|  | Fe | 18.24±1.31g | -11.8% | 22.02±1.33fg | -12.0% | 8.92±0.32fg | -7.6% |
|  | N+Fe | 18.70±1.33g | -9.7% | 22.69±1.48f | -9.4% | 9.05±0.27fg | -6.2% |
| OE-9 | H<sub>2</sub>O | 24.80±1.05c | 19.8% | 29.98±1.27c | 19.7% | 11.21±0.22bc | 16.1% |
|  | N | 26.28±1.00b | 26.9% | 31.64±1.64b | 26.4% | 11.70±0.29b | 21.2% |
|  | Fe | 26.04±0.78b | 25.3% | 31.58±1.14b | 26.1% | 11.57±0.31b | 19.8% |
|  | N+Fe | 27.74±1.29a | 34.0% | 33.74±1.56a | 34.8% | 12.58±0.22a | 30.3% |

Grain yield per plant, nitrogen use efficiency (NUE) and tiller numbers per plant of rice WT, *nlp4-1*, and OE-9 under different spraying conditions at the tillering stage in Bengbu in 2020 (May to October). Urea was used as the N source at 185 kg N/ha. The plants were transplanted in 8 rows × 10 plants for each plot (3.6 m<sup>2</sup>), and four replicates were used for each treatment. The experiment comprised four spraying conditions: i) H<sub>2</sub>O; ii) 0.5% urea (N); iii) 1 mM Fe(II)-EDTA (Fe); and iv) 1 mM Fe(II)-EDTA + 0.5% urea (N + Fe). At the early tillering stage, each plant was sprayed with 10 mL of a foliar solution 3 times at an interval of 5 days. Values are the mean ± SD (n=4 replicates, 60 plants per replicate for rice). Different letters above bars denote significant differences (P < 0.05) from Duncan's multiple range tests.

**Table S2.**
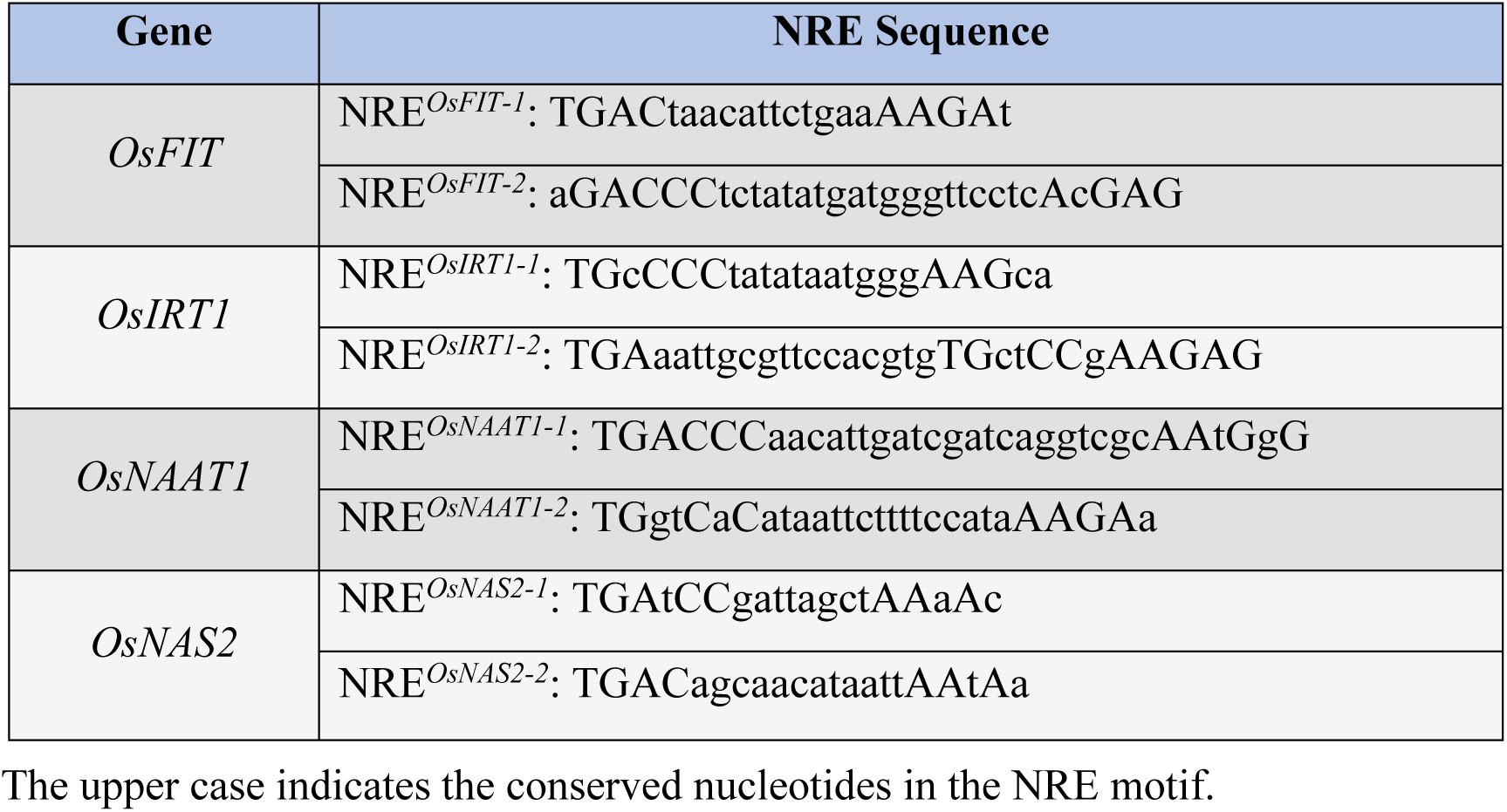
NREs found in the promoters of Fe-related genes targeted by OsNLP4.

**Table S3.**
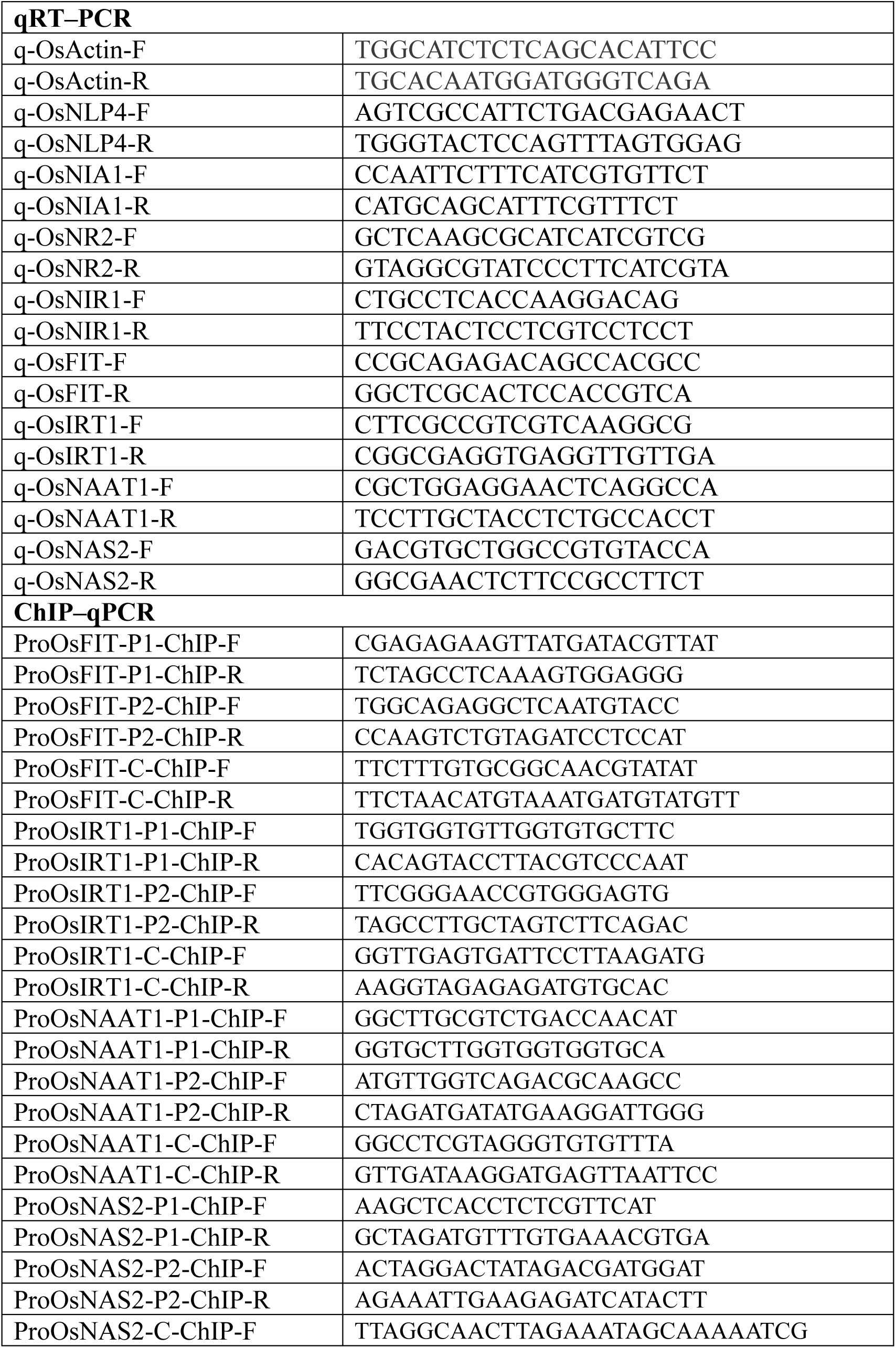
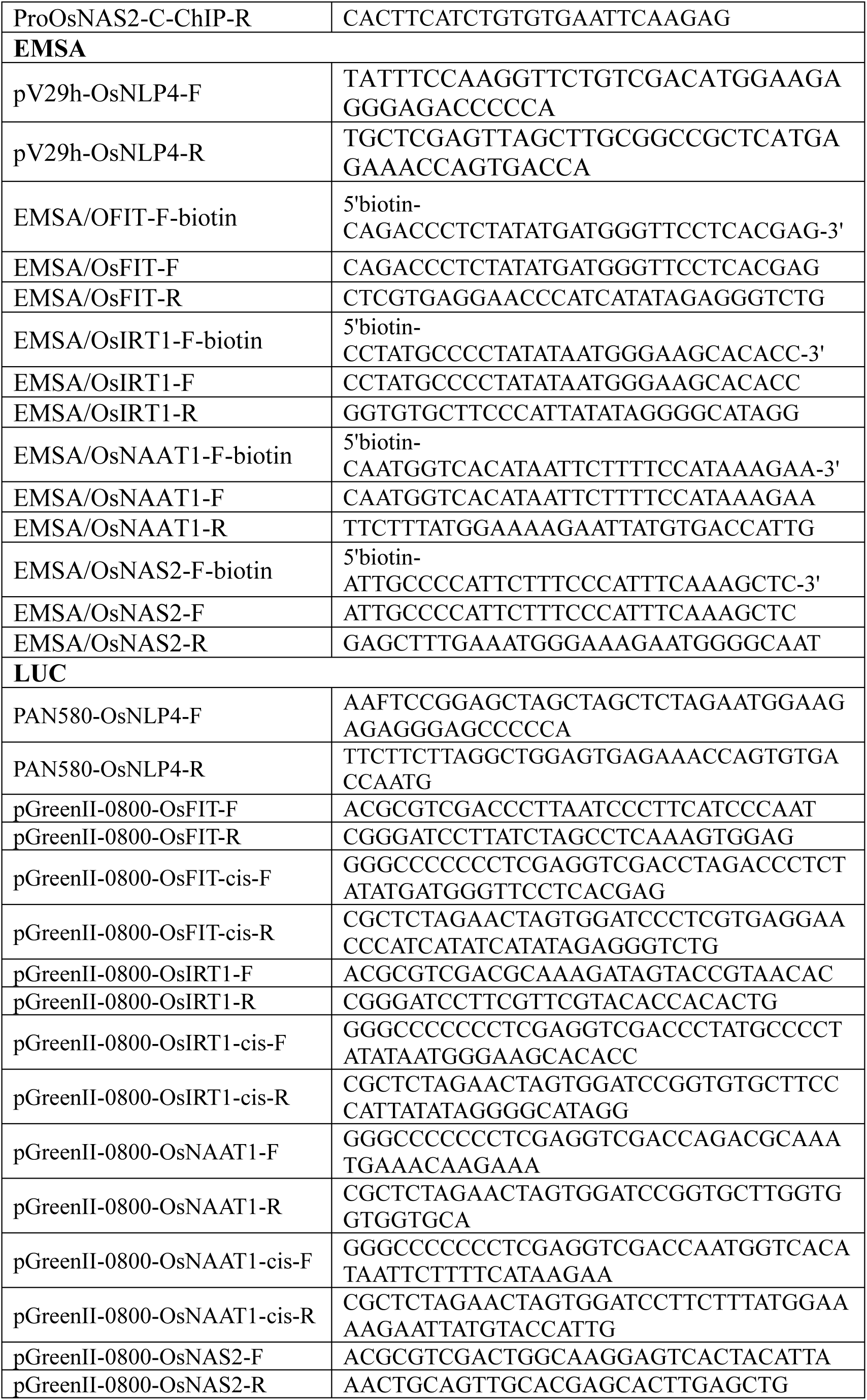
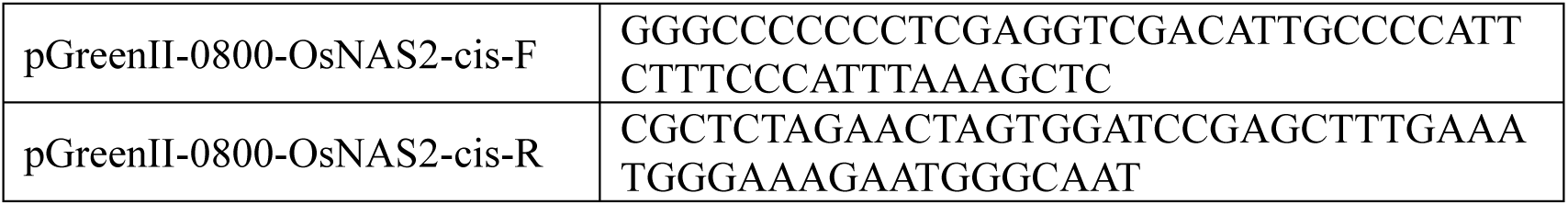
Primers used in this paper.

